# Cancer Histone Mutations Impact Binding and DNA Repair Processes, Leading to Increased Mutagenesis

**DOI:** 10.1101/2024.12.21.629746

**Authors:** Daniel Espiritu, Yiru Sheng, Yunhui Peng, Daria Ostroverkhova, Shuxiang Li, David Landsman, Maria Aristizabal, Anna R. Panchenko

## Abstract

Histones are key epigenetic factors for regulating the accessibility and compaction of eukaryotic genomes, affecting the replication, repair, and expression of DNA. Recent studies have demonstrated that histone missense mutations can perturb normal histone function, promoting the development of phenotypically distinguishable cancers. However, most histone mutations observed in cancer patients remain enigmatic in their potential to promote cancer development. To assess the oncogenic potential of histone missense mutations, we have gathered whole-exome sequencing data for the tumors of over 12,000 patients. Overall, histone mutations occurred in about 16% of cancer patients, although specific cancer types showed substantially higher rates. Using a combination of genomic, structural, and biophysical analyses, we found several predominant modes of action, where cancer missense mutations in histones affected acidic patches and protein binding interfaces in a cancer-specific manner and targeted interaction sites with specific DNA repair proteins. Consistent with this finding, we observed a high tumour mutational burden in patients with histone mutations affecting interactions of DNA repair proteins. We also identified potential cancer driver mutations in several histone genes, including histone H4-a highly conserved histone without previously documented driver mutations.

## Introduction

Histones are highly conserved nuclear proteins that are found ubiquitously in eukaryotic cells. They are key elements of nucleosomes, which consist of ∼147 base pairs of DNA complexed with two copies each of H2A, H2B, H3, and H4 histones^1,2^. Nucleosome core particles are flanked by linker DNA, where linker histone H1 can bind, forming chromatosomes. Nucleosomes and chromatosomes represent the fundamental subunits of chromatin. They are essential components for DNA packaging, act as hub points in epigenetic regulatory pathways, modulate DNA accessibility, and regulate DNA replication, repair, and transcription^1^. Chemical moieties (post-translational modifications, PTMs) can be added or removed from histone proteins, altering interactions with DNA and nucleosome-binding proteins^3^.

Although histones are evolutionarily conserved and critical for upholding genomic fidelity, histone genes are prone to accumulating mutations. Some missense mutations result in mutant histone proteins that can drive oncogenic phenotypes, also known as oncohistones^4,5^. There are several mechanisms whereby histones are thought to promote oncogenesis. Namely, histone mutations may locally perturb the reading, writing, and erasing of histone PTMs; disrupt histone-histone and histone-DNA interactions that are critical for forming stable nucleosomes; impact interfaces that are recognized by chromatin factors; or affect histone tail dynamics^6,7^. These local perturbations may scale up to global changes in chromatin dynamics, affecting DNA accessibility, repair, replication, and transcription^5^. Growing evidence suggests that histone mutations have a larger impact on cancer than previously thought^4,8,9^. While some mutations may be inconsequential (*i.e.* passenger mutations), others have been demonstrated to promote oncogenic phenotypes (*i.e.* driver mutations). This has led to several histone genes being recognized as cancer driver genes by the Catalogue of Somatic Mutations in Cancer (COSMIC) Cancer Gene Census (CGC)^10^. A handful of mutations (most notably, those in H1 and H3 histones) have been associated with clinically relevant outcomes such as overall survival^11^, tissue invasion^12,13^, and therapeutic response^14^. In recognition of H3 histone mutations’ impact on clinical prognosis, the World Health Organization (WHO) has expanded its classification of high-grade gliomas to include the H3.3 K27M mutation as a clinical biomarker^15,16^.

Histones are unique in that they are encoded by multiple genes^17^. Canonical histone genes are found in histone gene clusters, are deposited in a DNA replication-dependent manner, lack introns, and undergo unique 3′-end processing during transcription^18–20^. In contrast, variant histones are not found in clusters, their mRNA products can be expressed at any point in the cell cycle, their transcripts are polyadenylated, and they contain introns, allowing them to produce multiple transcripts and different protein isoforms.

While a handful of histone mutations have a demonstrated role in oncogenesis, there are many more observed in cancer with undetermined consequences^4,14,21^. Specifically, it has yet to be determined if these mutations meaningfully contribute to cancer development. Furthermore, mechanisms that allow oncohistone mutations to drive cancer development also remain obscure. To elucidate the role that histone mutations play in cancer *en-masse*, we have gathered a large whole exome sequencing (WES) dataset detailing somatic mutations for 11,950 cancer patients among 68 cancer types. Using various computational methods, we identified histone missense mutations that act as probable cancer drivers within specific cancer types, as well as in pan-cancer. Furthermore, we leveraged histone interaction networks to determine if missense mutations preferentially localize to histone interaction interfaces. To complement this analysis, we have also quantified how histone missense mutations may disrupt protein-protein and protein-DNA interactions using multiple methods (SAAMBE-3D, SAMPDI-3D, and MutaBind2)^22–24^. As disruption of histone PTM signaling is an emerging oncogenic mechanism, we identified additional mutations that affect histone PTM sites. Finally, we investigated how histone mutations may drive mutagenesis, given previous studies showing that histone mutations interfere with DNA repair^25,26^.

We found that the frequency of patients with predicted histone driver mutations depends on the cancer type, with malignant diffuse large B-cell lymphoma showing the largest frequency of predicted histone driver mutations. Furthermore, we reported that histone mutations were strongly enriched at acidic patch residues. In concordance with this, histone mutations preferentially aggregated at histone interfaces for specific proteins. We proposed that Tonsoku-Like protein (TONSL), Death Domain Associated Protein 6 (DAXX), FAcilitates Chromatin Transcription (FACT) complex subunit SSRP1, Histone Chaperone ASF1A, DNA Replication Licensing Factor MCM2, BRCA-1 Associated RING Domain Protein 1 (BARD1), Lysine-specific histone demethylase 1A (LSD1), and Histone acetyltransferase KAT6B may act as mediators of histone mutation-driver oncogenesis. Moreover, we noted that patients harboring histone mutations on interfaces for these proteins had a significantly higher tumor mutational burden than patients without them. Of particular interest, we identified several H4 histone mutations that could serve as probable drivers of oncogenesis, further noting that they affected interactions with possible oncogenic mediators. Among these, the H4 R67P mutation is highly disruptive of nucleosome structures and was predicted as a driver in multiple cancer types. In summary, our findings suggest that histone mutations drive oncogenesis in a cancer-specific manner by perturbing interactions with other proteins, including DNA repair proteins, and increasing tumor mutation rates.

## Results

### Histone missense mutations drive oncogenesis in a cancer-specific manner

Tumor WES and clinical data were gathered from the Genomic Data Commons for 11,950 patients, all of whom were adults over the age of 18 at the time of diagnosis, except for one individual. Among these patients, missense mutations represented the largest proportion (60%) of all observed histone mutation types, with 2,803 distinct histone missense mutations (referred to as “histone mutations” thereafter) spread across 2,029 patients (∼16% of all patients). Given their prevalence, we focused on histone missense mutations and refer to them as “histone mutations” throughout the text. To evaluate the novelty of our dataset, we compared it to an existing data set of cancer histone mutations derived from WES and targeted sequencing studies^4^. After removing targeted sequencing studies from the aforementioned dataset, we found that 39% of our patients with histone mutations and 38% of histone mutations were novel and not present in this previous data set^4^. In total, we identified 1065 new unique mutations.

To determine if our list of histone mutations includes novel drivers of oncogenesis, we predicted cancer driver status using MutaGene^27,28^. Because MutaGene predicts driver status based on the observed mutation frequency, samples with high tumor mutational burdens (TMB) were removed for this analysis (see Methods). This was done as the abnormally high TMBs in these tumors may be explained by factors other than selective pressures acting in the broader population. Driver status prediction was conducted using a pan-cancer model and for specific cancer types by separating mutations based on patient primary diagnosis (68 in total, Methods). Only one mutation, *CENPA* Y109H, was predicted as a driver of oncogenesis in pan-cancer, but it only occurred in two patients. Repeating this analysis by grouping samples based on primary cancer diagnosis revealed 218 distinct cancer-type specific driver mutations in histones across 55 cancer types (Supplementary Table 1). Importantly, most mutations were predicted as drivers in a single cancer type, highlighting the cancer-specific nature of histone cancer driver mutations. Although most predicted cancer drivers were rare mutations detected in a single patient, 29 driver mutations were recurrent and observed in more than one patient. Six mutations were predicted as drivers in multiple cancer types. Among them, the H3 K27M, G34R, and K36M mutations have been well documented previously^5,8,9,29,30^, whereas *H2BC21* F70L was less well-documented^31^, but shown to increase H2A/H2B dimer exchange via Nap1^32^, which may be related to its destabilizing effect on the nucleosome (mean SAAMBE-3D ΔΔG 1.53 kcal/mol, mean MutaBind2 ΔΔG = 1.35 kcal/mol). The *H3C2* E133Q mutation was predicted as a driver in mixed glioma and diffuse large B-cell malignant lymphoma. Importantly, for the first time we predicted a driver mutation in H4 histone type, *H4C3* R67P, in multiple cancer types. It was also highly disruptive of nucleosomal histone-histone interactions (mean SAAMBE-3D ΔΔG = 2.64 kcal/mol; MutaBind2 on representative PDB structures 3X1T and 7A08, mean ΔΔG = 2.14 kcal/mol) and overlapped DAXX, MCM2, and BARD1 interaction interfaces. As will be discussed in next sections, histone driver mutations are enriched at interaction interfaces for these partners and patients with mutations at interfaces for these partners exhibit a high TMB.

Next, we examined if the frequency of all histone mutations and histone driver mutations was dependent on the cancer type (Supplementary Tables 2,3). We found significant depletions (Fisher’s Exact, q < 0.05, Odds Ratio (OR) < 1) of all and driver mutations in adult glioblastoma (pediatric cancers were not represented in this study), infiltrating duct carcinoma, and papillary carcinoma. By contrast, significant enrichments (OR > 1) of all and driver mutations were detected in diffuse large B-cell malignant lymphoma and transitional cell carcinoma (Fig. 1A). The cancer type with the strongest enrichment for mutations was diffuse large B-cell malignant lymphoma (OR for driver = 43.3 and OR for all mutations = 7.98), where 53% of patients harbored a histone driver mutation. Within this cancer type, driver mutations were found across all histone types and were most abundant when all histone H1 proteins were considered collectively (Fig. 1B). Focusing on specific histone proteins, we found that cancer drivers in H4, H1.4, and H1.2 were the most frequent (Fig. 1C).

**Figure 1.**
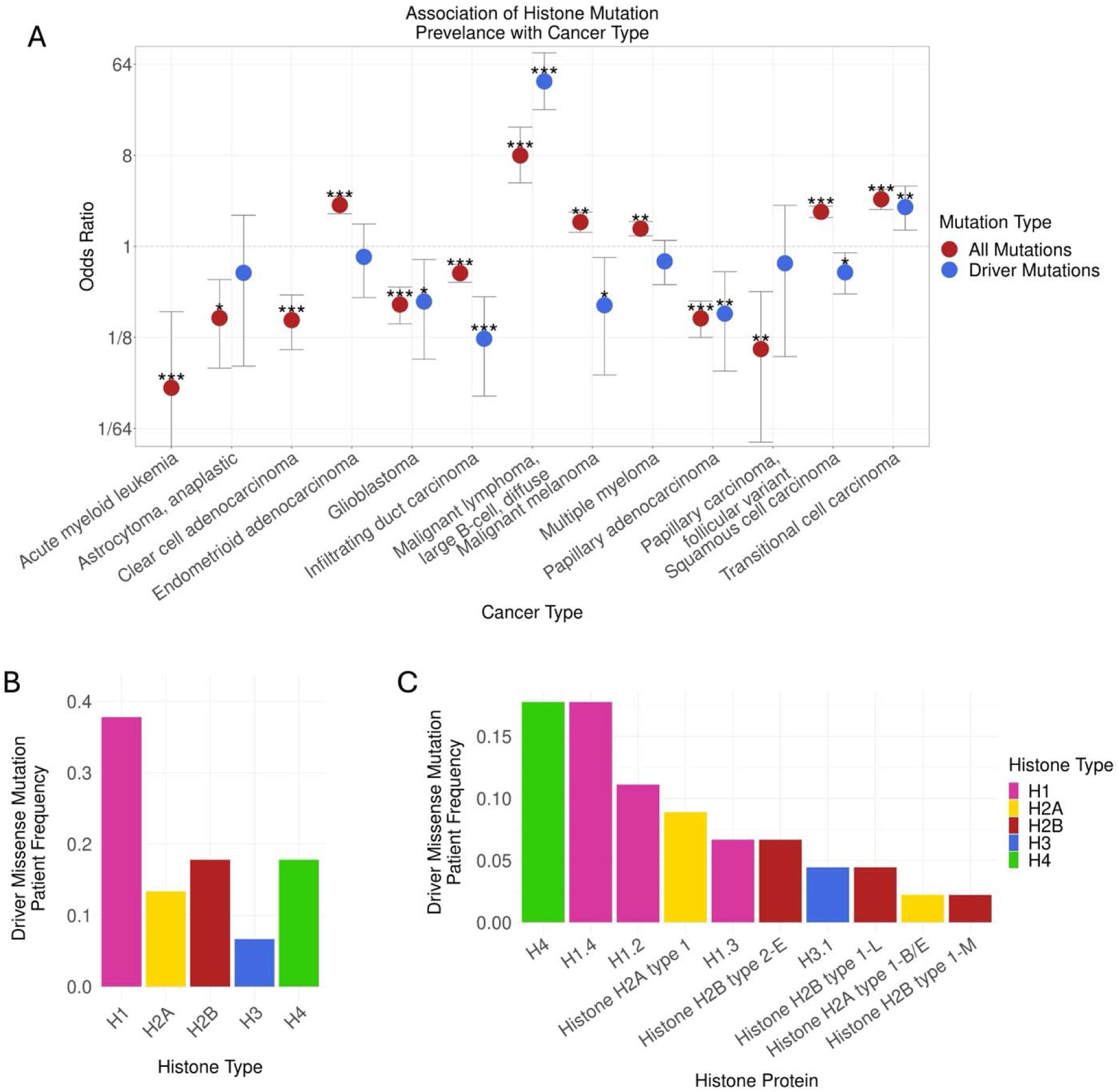
Histone missense mutations across cancer types, histone types, and proteins. A) Cancer types found to have a significantly high or low number of patients with at least one histone mutation (red) or cancer-specific histone driver mutation (blue), when compared to all patients not within the given cancer type (Fisher’s Exact, q < 0.05, Benjamini-Hochberg, FDR = 0.05). B) The frequency of patients with a cancer-specific histone missense driver mutation in a given histone type are shown for Diffuse Large B-cell Malignant Lymphoma. C) The frequency of patients with histone driver mutations in a gene encoding a given histone protein are shown for Diffuse Large B-cell Malignant Lymphoma. Frequencies in B-C) were calculated as the number of Diffuse Large B-cell Malignant Lymphoma patients with histone driver mutations in a gene for a given histone type (B) or protein (C), divided by the total number of Diffuse Large B-cell Malignant Lymphoma patients after removing tumor mutational burden outliers. * - q < 0.05, ** - q < 0.01, *** - q < 0.001

### Histone mutations co-localize with specific types of interaction interfaces, acidic patch residues, and PTM sites

Previous studies found that many histone mutations co-localize with histone interaction interfaces^4^, but whether these associations are statistically significant has not been explored. Powered by our recently constructed human histone interactome^33^, we asked if there are statistically significant associations between the occurrence of histone mutations and nucleosomal histone-histone interfaces, nucleosomal histone-DNA interfaces, histone-non-histone protein (histone-NHP) interfaces, acidic patch residues, or PTM sites (see Methods for definitions). To identify common themes across cancer types, we performed this analysis using the pan-cancer model, where all mutations were considered collectively or separated by whether they were recurrent (occurring in more than one patient) or non-recurrent. Predicted driver status was not considered in this analysis because only one mutation was predicted to be a driver in pan-cancer. Furthermore, associations were tested for both empirically derived and inferred interaction interfaces (see Methods). The associations reported below were consistent regardless of the type of statistical test performed (Binomial or Fisher’s exact) or the type of data supporting the interaction (empirically derived or inferred) unless otherwise specified (Fig. 2A, B; Supplementary Figure 1, 2, 3; Supplementary Tables 4-13).

**Figure 2.**
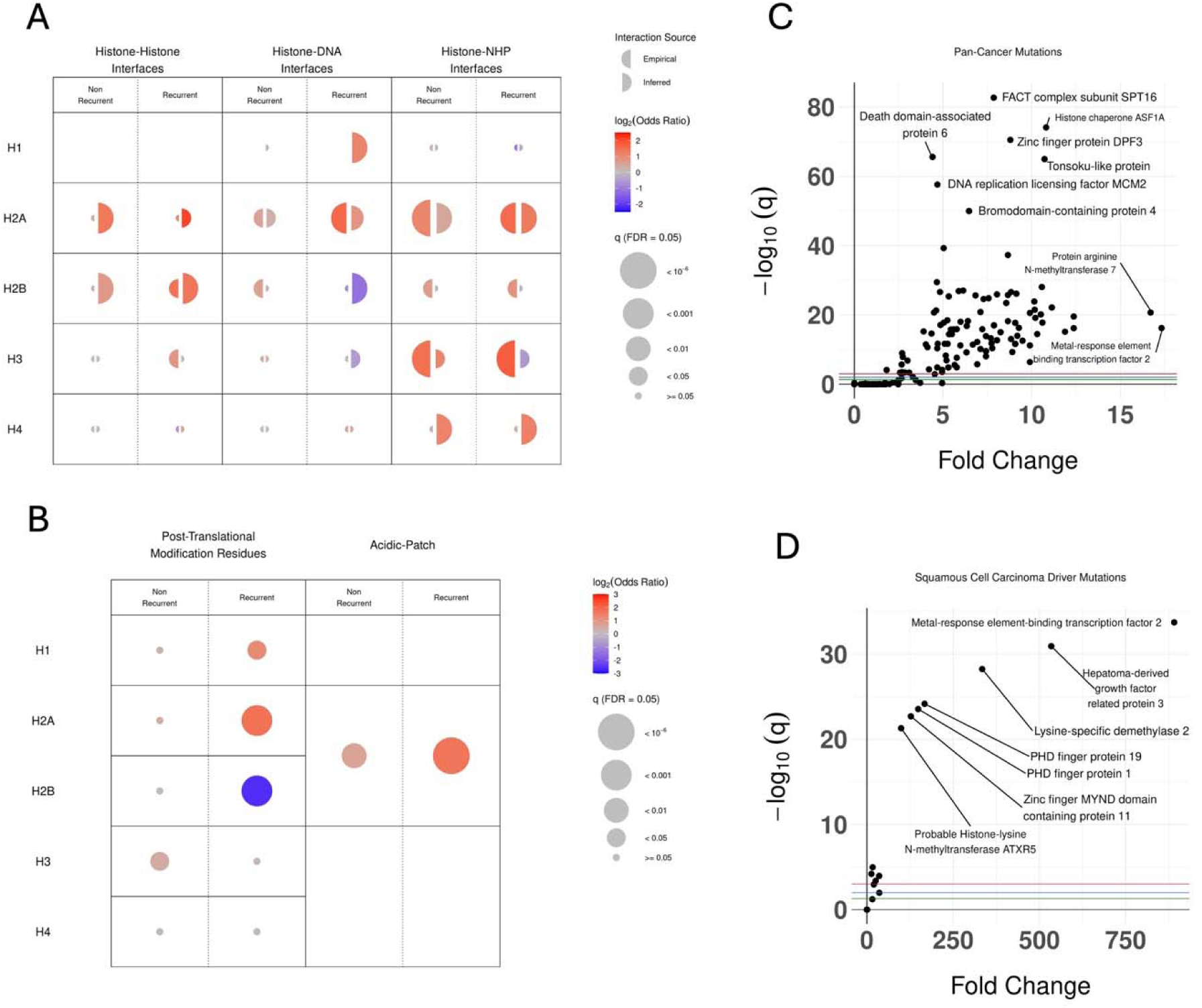
Histone mutations occur at functionally important residues and interfaces with specific interaction partners. A) Association of histone mutations in pan-cancer with empirically derived (left semi-circle) and inferred (right semi-circle) interaction interfaces usinf Fisher’s exact test. Results are stratified by histone type and mutation recurrence. Recurrent mutations are those found in more than one patient. B) Association of histone mutations in pan-cancer with PTM sites and acidic patch residues. C) and D) Volcano plots showing q-values and effect sizes (fold-change) for the association of histone mutation occurrence with interfacial residues for specific non-histone proteins (NHPs) for all histone mutations in pan-cancer (C) and driver mutations in Squamous Cell Carcinoma (D). q-value thresholds corresponding to 0.05 (green), 0.01 (blue), and 0.001 (red) are shown. Fold-change was calculated as the number of observed mutations at an interface for a given partner, divided by the number of expected mutations. Partners with the lowest q-values and highest fold-changes in C-D) have been labelled. q-values were adjusted using the Benjamini-Hochberg method (FDR = 0.05).

Several major patterns were detected between histone mutations and interaction interfaces (Fig. 2A, Supplementary Fig. 1A). Namely, significant enrichments were found for both recurrent and non-recurrent H2A mutations with DNA and NHP interfaces. For H2B histones, significant enrichments were found between recurrent mutations and histone-histone interfaces. The association between histone mutations and NHP binding partners was more variable for H3 histones. Specifically, recurrent and non-recurrent mutations were significantly enriched on empirically derived NHP interfaces, but this was not seen for recurrent mutations when inferred interactions were considered. Although the limited representation of H1 proteins in the nucleosomal interactome prevented us from testing empirical H1-DNA interfaces, we did find that recurrent H1 mutations were significantly enriched at inferred H1-DNA interfaces. Finally, histone H4 stands out as having the least number of significant associations.

Strong significant enrichments were found for both recurrent and non-recurrent mutations on acidic patch residues (Fig. 2B, Supplementary Fig. 1B). When compared to non-acidic patch interfacial residues, rather than all residues, this significant enrichment persisted for recurrent mutations (Supplementary Fig. 1B, 2). As for PTM sites, significant enrichments were found with recurrent mutations in H1 and H2A histones, and non-recurrent mutations in H3 histones, although with a weaker signal. Conversely, a strong depletion was observed between recurrent mutations and PTM sites for H2B histones. Results concerning all mutations, regardless of recurrence, can be seen in Supplementary Fig. 3 and Supplementary Tables 4-13.

Of the three categories of histone-binding partners (histone, DNA, NHP), the NHP category is the most structurally and functionally diverse. To identify potential mechanisms by which histone mutations may drive oncogenesis, we examined the localization of histone mutations with respect to interfaces for all 366 NHP binding partners individually. Overall, we found that interfaces for 133 partners were significantly enriched for histone mutations in pan-cancer (Binomial and Fisher’s exact test q < 0.05, Fig. 2C). To determine if histone mutations associate with interfaces for specific NHPs in specific cancer types, we repeated this analysis by examining each cancer type separately (Supplementary Tables 14, 15) and using only predicted driver mutations (Supplementary Tables 16, 17). Overall, we found that driver mutations were enriched at interfaces with 53 different NHPs within 11 cancer types. For 24 out of these 53 NHPs, the histone driver mutations occurred within H4 histones, an interesting finding given that driver mutations on H4 have not been documented previously (Supplementary Table 18). Among these NHPs are DAXX, MCM2, ASF1A, TONSL, SSRP1, and BARD1. We also noted significant enrichments for squamous cell carcinoma driver mutations at interfaces with Polycomb-like proteins Metal-Response Element-Binding Transcription Factor 2 (MTF2), PHD Finger Protein 19 (PHF19), and PHD Finger Protein 1 (PHF1) (Fig. 2D). Further analysis revealed that all driver mutations on interfaces for these Polycomb-like proteins were H3 K36M mutations in head and neck squamous cell carcinoma (HNSCC) patients, a mutation previously shown to drive cancer development in HNSSC patients by disrupting Polycomb Repressive Complex 2 function, which Polycomb-like proteins regulate^29,34^.

### Effects of histone mutations on binding affinity

Prompted by our finding that histone mutations significantly associate with specific binding interfaces, we examined if histone mutations lead to changes in binding affinity (ΔΔG). To systematically explore this possibility, we used SAAMBE-3D and SAMPDI-3D to estimate the effects of mutations on histone-protein (Supplementary Table 19) and histone-DNA interactions (Supplementary Table 20), respectively. In total, we obtained ∼190,000 ΔΔG estimates for 1,370 mutations, using all available histone structures (536 structural evidences).

To identify histone mutations that are highly disruptive of nucleosomal interactions, relative to other mutations, we calculated the change in binding affinity of histone-histone and histone-DNA interactions for all mutations in our dataset. All changes in binding affinity were converted to modified Z-scores and mutations were classified as highly disruptive outliers, either destabilizing (modified Z-score > 3.5) or stabilizing (modified Z-score < -3.5). While no mutations were found to be nucleosomal histone-DNA binding outliers, ten mutations were found to be nucleosomal histone-histone binding outliers (Fig. 3). Notably, seven of these mutations were predicted to be drivers or potential drivers in at least one cancer type, while the remaining mutations (H2A type 1 R81P, H2A type 2-C R81P, and H3.1 A111D) were not ranked by MutaGene as they occurred in patients removed after TMB filtering (see Methods). Of these, one mutation, H3.1 A111D, was found to stabilize H3.1-histone interactions (mean ΔΔG = -1.04 kcal/mol) while the rest were predicted to destabilize interactions. Of note, the H3.1 R131Q mutation was predicted to be highly disruptive of H3.1-histone interactions (mean SAAMBE-3D ΔΔG = 2.69 kcal/mol). This mutation is located near the *H3C2* E133Q mutation (also encoding H3.1), which was predicted to act as an oncogenic driver in multiple cancer types.

**Figure 3.**
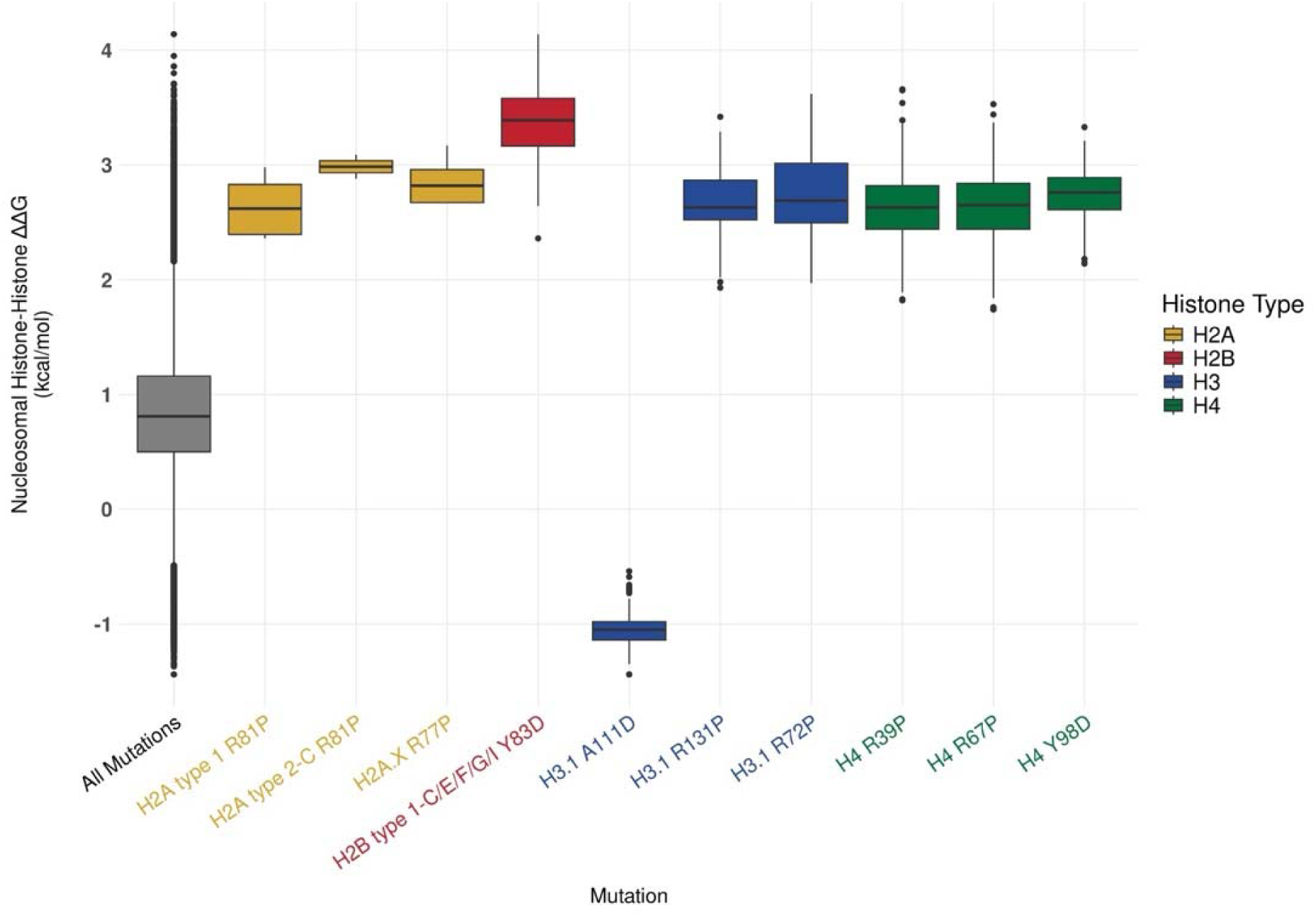
Histone mutations predicted as oncogenic drivers in multiple cancer types and highly disruptive of histone-histone interactions. Distribution of nucleosomal SAAMBE-3D ΔΔG prediction for histone mutations that were found to be hyper-disruptive outliers of nucleosomal histone-histone interactions. SAAMBE-3D ΔΔG values above zero indicate a destabilization of interactions, whereas values below zero indicate a stabilization of interactions. Mutations and ΔΔG distributions have been color-coded by the histone type they occur in.

Having identified enrichments between the presence of histone mutations and interfaces with histone-binding proteins (Fig. 2), we examined if mutations can differentially alter specific histone-protein binding affinities. Comparing the distributions of changes in binding affinity for a given protein binding partner (histone or NHP) to the cumulative distribution of binding affinity changes of all other histones and NHPs revealed 32 specific binding partners predicted to be disproportionally affected by cancer histone mutations (Mann-Whitney U test, q < 0.05) (Supplementary Table 21). Notably, mutations on interfaces with ASF1A, TONSL, and the heterodimeric FACT complex^35^ subunits SSRP1 and SPT16 were found to be more destabilizing as determined by significantly higher ΔΔG values when compared to mutations on interfaces for other proteins. When looking within specific cancer types we found that binding with TONSL, Histone H2B Type 1-J, Histone H2B type 1-C/E/F/G/I, Chromobox Protein Homolog 1, Peregrin, and SPT16 was disproportionately disrupted by histone mutations. Consistent with our results showing associations between histone mutations and NHP interfaces, mutations affecting TONSL interactions were found to be more destabilizing in adenocarcinoma, endometrioid adenocarcinoma, and squamous cell carcinoma, when compared to mutations affecting interactions for other proteins.

### The presence of histone mutations is associated with high tumor mutational burden

Given the link between histone mutations and proteins involved in DNA damage repair (e.g. TONSL and DAXX), we examined if the presence of histone mutations and their NHP interfacial status was associated with the number of mutations per patient (Tumor Mutational Burden, TMB). Kruskal Wallis and post-hoc Dunn tests showed that patients with histone mutations (both interfacial and non-interfacial) had a significantly higher TMB than patients without histone mutations in pan-cancer (Fig. 4A) and cancer types examined individually (q < 0.05, rank-biserial correlation coefficient ranges from 0.16 to 0.86, Supplementary Tables 22, 23). Furthermore, significantly higher TMB was found for patients with interfacial histone mutations when compared to patients with non-interfacial histone mutations in two cancer types: serous cystadenocarcinoma (rank-biserial correlation coefficient = 0.35) and endometrioid adenocarcinoma (rank-biserial correlation coefficient = 0.28).

**Figure 4.**
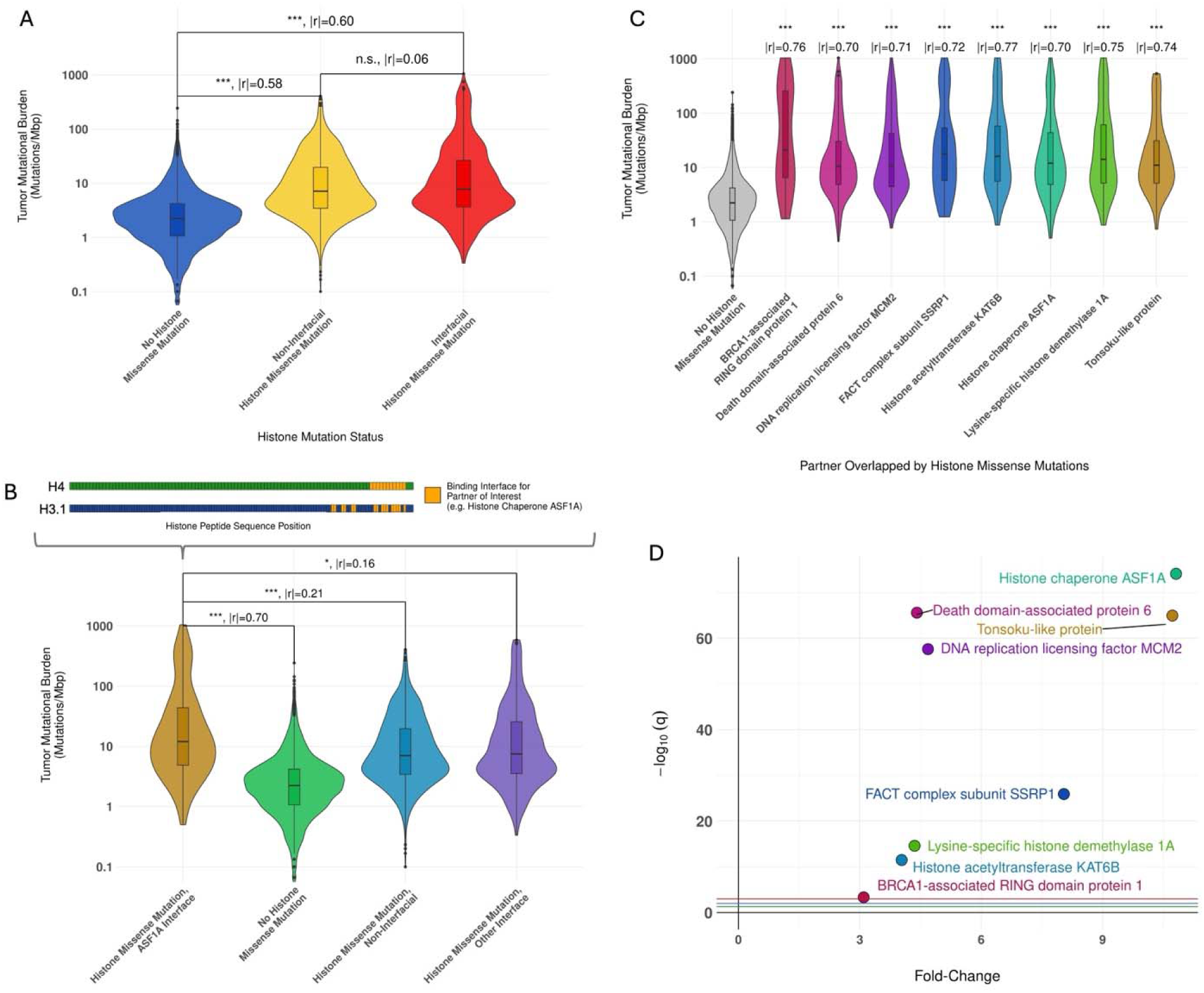
Histone mutations as promoters of mutagenesis in pan-cancer. A) Pairwise comparisons of tumor mutational burden (TMB) distributions in patients grouped by histone mutation status (Dunn test; absolute rank-biserial correlation coefficient, |r|). B) Schematic showing how non-histone proteins (NHPs) were identified as potential mediators of histone mutation-associated TMB. Patients with histone mutations on interfaces with an NHP of interest were grouped (top) and their TMB distributions were examined (bottom). The TMB distributions were compared to the TMB distribution seen in patients without histone mutations, with mutations at sites that are not involved in physical interactions, and with mutations at sites that contact other NHPs, using Mann-Whitney U tests. C) Comparison of the TMB distributions for patients with histone mutations that overlap the indicated NHP compared to the distribution seen in patients without histone mutations (Mann-Whitney U, rank-biserial correlation). D) Volcano plot showing the enrichment of histone mutation occurrences at histone binding interfaces for NHPs shown in C) (Binomial test). q-value thresholds correspond to 0.05 (green), 0.01 (blue), and 0.001 (red). NHPs have been labelled and color coded to match C). Fold-change was calculated as the observed number of histone mutations at a given NHP interface, divided by the expected number of mutations. All reported q-values were obtained via the Benjamini-Hochberg method (FDR = 0.05). n.s. - q > 0.05, * - q < 0.05, ** - q < 0.01, *** - q < 0.001

To investigate the mechanisms by which histone mutations may increase TMB, we examined if patients with mutations overlapping specific NHP interfaces have a significantly higher TMB compared to patients without histone mutations, patients with histone mutations not on NHP interfaces, and patients with histone mutations at interfaces for other NHPs (Mann-Whitney U, q < 0.05, see Methods, Fig. 4B). This analysis identified MCM2, DAXX, LSD1, SSRP1, KAT6B, TONSL, ASF1A, and BARD1 as binding partners in pan-cancer that may link histone mutations to high TMB (Fig. 4C, Supplementary Tables 24). Consistent with this finding, we note that pan-cancer histone mutations were significantly enriched at interfaces with all these binding partners (Fig. 4D). In addition, pan-cancer histone mutations affecting histone interactions with SSRP1, KAT6B, ASF1A, and TONSL had significantly higher ΔΔG values when compared to other proteins. In the context of specific cancer types, DAXX, SPT16, MCM2, ASF1A, and TONSL interfacial histone mutations were also found to be associated with high TMB in more than one cancer type (Supplementary Table 24). Mann-Whitney U tests using permutations to generate a null distribution were conducted to obtain second estimates of p-values showed similar results (Supplementary Table 25).

Previously, we showed a positive link between the presence of histone and *POLE* mutations^21^. Mutations in *POLE*, which encodes DNA polymerase ε, are associated with extremely high TMB, especially in serous cystadenocarcinoma and endometrioid adenocarcinoma^36,37^. To ensure that our results were not biased by *POLE* mutant status, we repeated our analyses after removing patients with *POLE* mutations and found that the significant differences in TMB between patients with and without histone mutations persisted (rank-biserial correlation coefficient ranges from 0.14 to 0.67, Supplementary Tables 26, 27 and Supplementary Fig. 4). Interestingly, histone mutations that overlapped TONSL and DAXX interfaces were associated with high TMB, independent of *POLE* mutations (Supplementary Tables 28, 29).

### Newly discovered H4 and H3.3 mutations may drive oncogenesis by disrupting TONSL and DAXX interactions, important DNA repair proteins

Our results so far suggest that histone mutations may drive oncogenesis by affecting TONSL function. TONSL is a protein that binds newly synthesized H4 proteins and assists in double-stranded break repair^38^. Our analyses showed that histone mutations were enriched at TONSL interfaces in several cancer types with very high effect sizes (fold-change ≥ 10, Fig. 5A) and are predicted to destabilize TONSL binding (Fig. 5B). Moreover, histone mutations on TONSL interfaces associated with high TMB in pan-cancer, an effect independent of *POLE* mutations (Fig. 5C). The TONSL ankyrin-repeat domain (ARD) contacts 22 residues on H4 (Fig. 5D) and two residues on H3.3^38^. While numerous H4 cancer mutations were found on TONSL interfaces, only one on H3.3 mutation was observed. Mutations in both the H4 tail (Fig. 5E, ΔΔG from 0.24 to 2.98 kcal/mol) and H4 core domain (Fig. 5F, ΔΔG from 0.25 to 1.78 kcal/mol) show a trend towards highly destabilizing interactions with the TONSL ARD. Predictions using MutaBind2 show comparable results (Supplementary Table 30).

**Figure 5.**
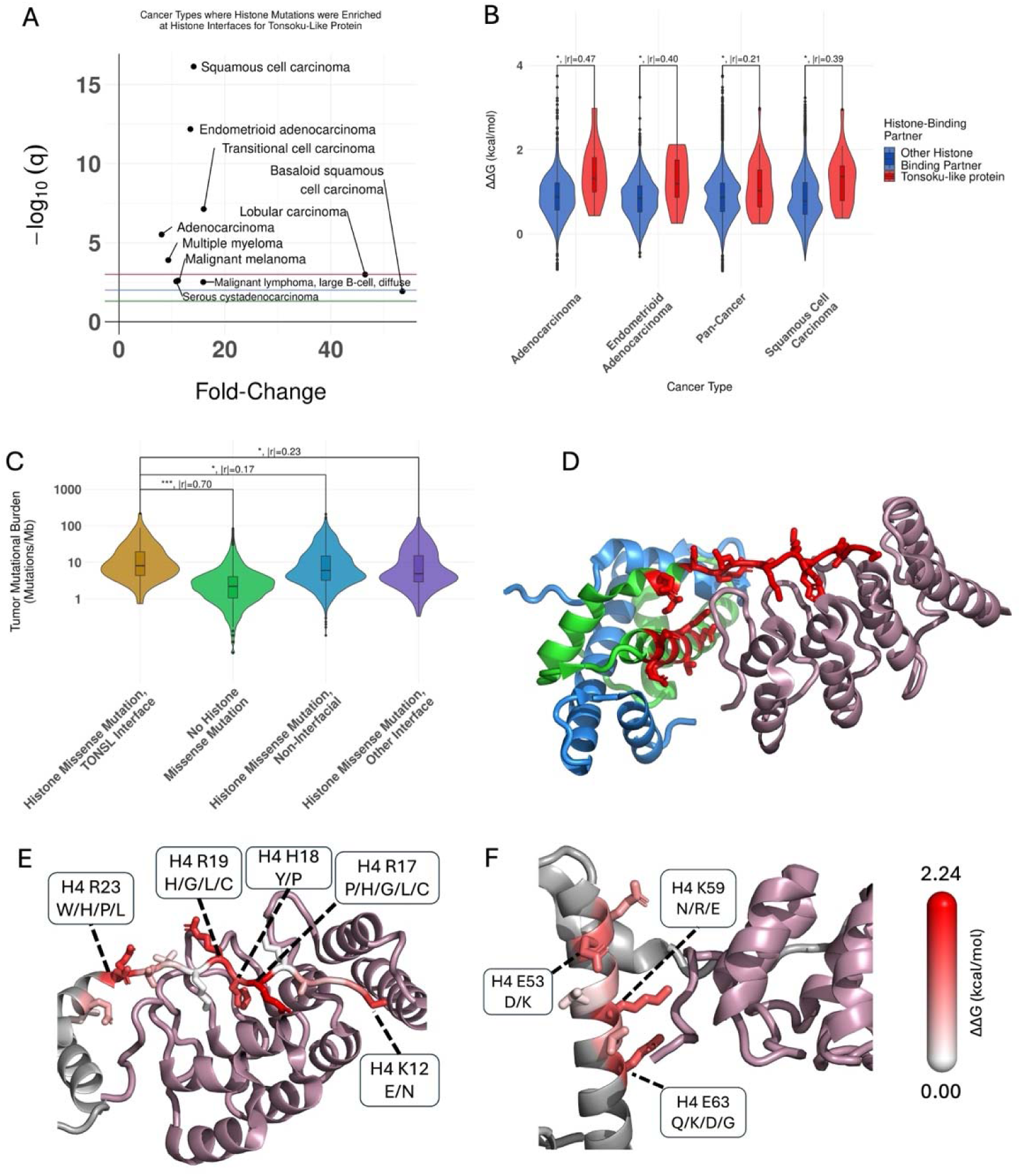
Tonsoku-Like Protein may mediate elevated TMB associated with histone mutations. A) Cancer types where histone mutations were found to be significantly enriched (Binomial test, q < 0.05) at histone interfaces with TONSL. Fold-change was calculated as the number of observed mutations at a TONSL interface divided by the number of expected mutation observations. q-value thresholds correspond to 0.05 (green), 0.01 (blue), and 0.001 (red). B) The ΔΔG distributions of histone mutations affecting TONSL interactions, compared to the ΔΔG distributions of histone mutations affecting interactions for other partners. Results are separated by cancer type (Mann-Whitney U; absolute rank-biserial correlation coefficient, |r|). C) Pairwise comparison of TMB distributions for patients harboring histone mutations on TONSL interfaces vs the indicated patient groups (Mann-Whitney U). All reported q-values in A-C) were obtained via the Benjamini-Hochberg method (FDR = 0.05). D) Structure of the H3.1-H4 dimer (blue-green, respectively) in complex with TONSL (purple). Sites of H4-TONSL interaction are colored red. Structure showing mean ΔΔG for H4 tail domain mutations E) and core domain mutation F) mapped to TONSL interfaces. ΔΔG changes are represented using a white-red color gradient (low-high, respectively). H4 residues that are not interfacial for TONSL are colored gray, alongside interfacial core domain residues in E) and tail residues in F). Side chains are shown for wild-type residues at TONSL interaction sites. Patients without *POLE* mutations were considered in C-F). Figures D, E, and F are based on PDB: 5JA4. * - q < 0.05, ** - q < 0.01, *** - q < 0.001

Akin to TONSL, DAXX function may also be commonly impacted by histone mutations. DAXX acts as a histone H3.3-H4 chaperone during double-stranded break repair^39^. Not only were histone mutations enriched at histone-DAXX interfaces in several cancer types (fold-change ≥ 3.47, Fig. 6A), but these mutations were also associated with a high TMB (Fig. 6B). DAXX contacts 42 residues on H4 and 77 residues on H3.3. Only one driver mutation was found at an H3.3-DAXX interface in transitional cell carcinoma, whereas 18 distinct driver mutations were found at H4-DAXX interfaces, among 31 patients. Generally, mutations were observed at a lower frequency on H3.3-DAXX interfaces (Fig. 6C) compared to H4-DAXX interfaces (Fig. 6D, Supplementary Fig. 5). Interestingly, patients with H3.3-DAXX interfacial mutations had a significantly higher TMB (median = 14.0 mutations/Mb) than patients with H4-DAXX interfacial mutations (median = 7.1 mutations/Mb, Mann-Whitney U, rank-biserial = 0.34, p < 0.05). Moreover, H3.3-DAXX interfacial mutations had significantly higher ΔΔG values (median = 1.01 kcal/mol) when compared to H4-DAXX interfacial mutations (median = 0.77 kcal/mol, Mann-Whitney U, rank-biserial = 0.23, p < 0.05), although these predictions were not validated using MutaBind2 (Supplementary Table 30).

**Figure 6.**
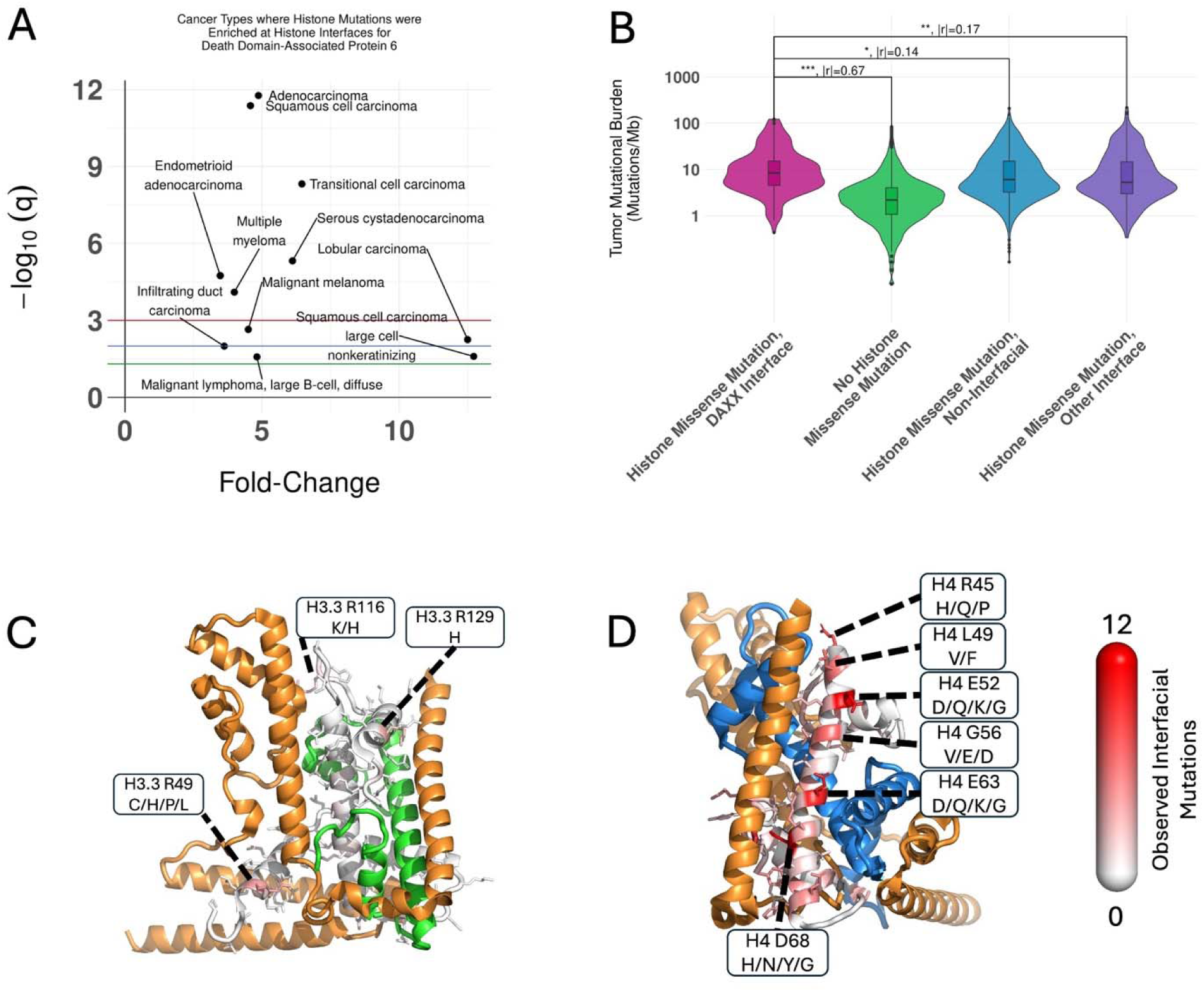
Death Domain Associated Protein 6 may be involved in the increased TMB associated with histone mutation. A) Cancer types where histone missense mutations were found to be significantly enriched (Binomial test, q < 0.05) at histone interfaces with DAXX. Fold-change was calculated as the number of observed mutations at a DAXX interface, divided by the number of expected mutation observations. q-value thresholds correspond to 0.05 (green), 0.01 (blue), and red (0.001). B) Pairwise comparison of TMB distributions of patients harboring histone mutations on DAXX interfaces and the indicated patient groups (Mann-Whitney U). C-F) An H3.3-H4 dimer (blue-green, respectively) in complex with DAXX (orange) (PDB: 4HGA). The number of observed histone mutations on DAXX-interfacial residues is shown by a white-red color gradient (low-high, respectively). In C) the color gradient is applied to H3 residues, with non-interfacial H3 residues also colored white. In D-F), the color gradient is applied to H4 residues, with non-interfacial H4 residues also colored white. Side chains are shown for interfacial wild-type residues. All reported q-values in A-B) were obtained via the Benjamini-Hochberg method (FDR = 0.05). Pan-cancer patients without *POLE* mutations were considered in B-F). * - q < 0.05, ** - q < 0.01, *** - q < 0.001

**Figure 7.**
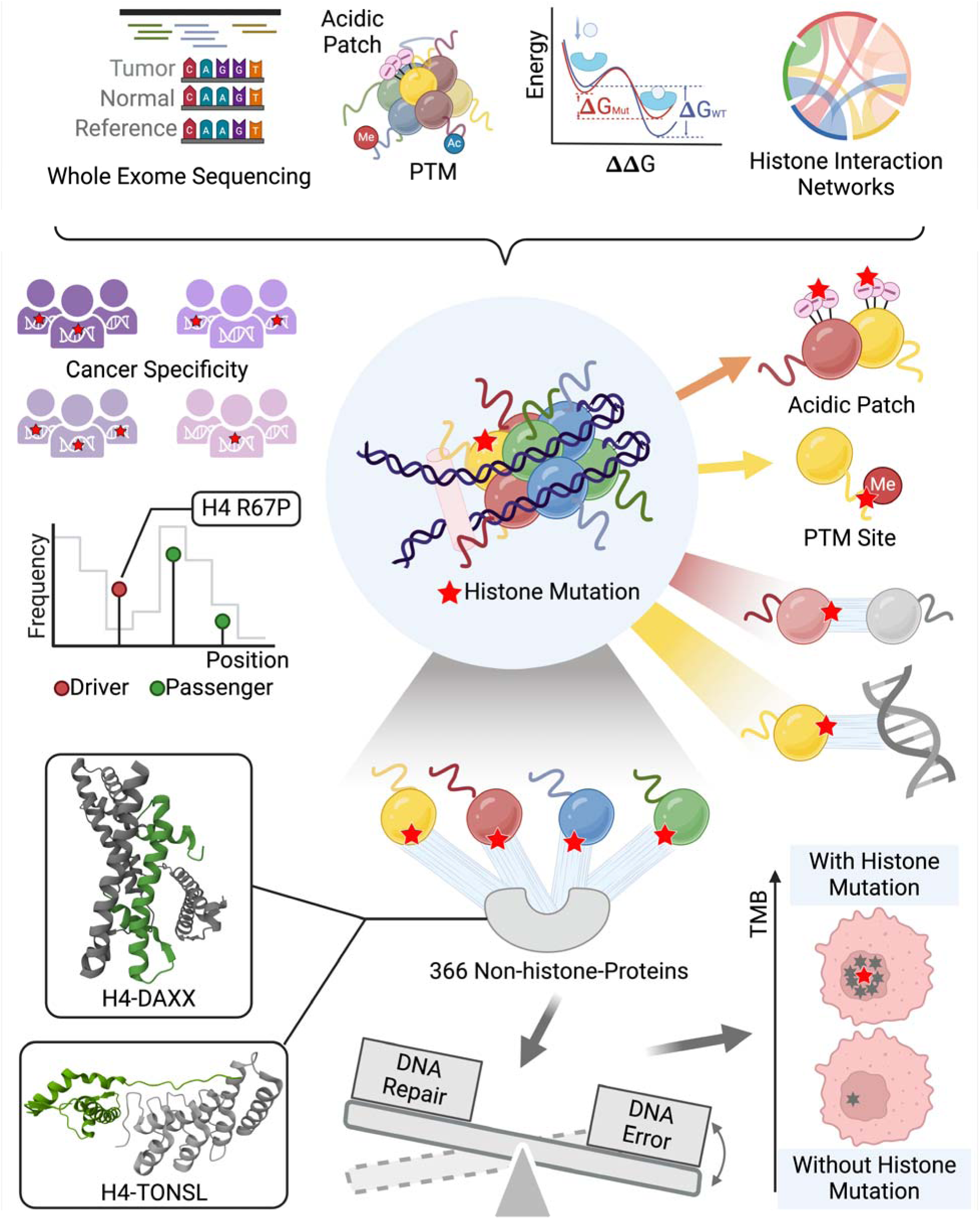
A summary of the main findings of the paper. Created in BioRender. Sheng, Y. (2024) https://BioRender.com/j40m779

## Discussion

Dysregulated epigenetic states are hallmarks of cancer and often arise from genetic alterations in epigenetic regulators, including histones. In this work, we examined the role that histone mutations play in oncogenesis within the context of pan-cancer and specific cancer types (please see a summary Figure 6). While some histone mutations are capable of driving cancer development, there are numerous other mutations observed in cancer with unknown consequences^4^. As the presence of histone mutations may be associated with patient prognosis, determining if, and how, they are likely to contribute to cancer development is necessary to develop biomarkers or therapeutic interventions^11,40–42^. By bringing together complementary genomic, structural, and biophysical data, we identify histone mutations that may drive cancer development and the mechanisms by which they exert their effects.

Our findings indicate that most histone mutations operate in a manner specific to each cancer type, aligning closely with the cell-type specificity inherent to epigenetic regulation. We have identified several cancer types in which histone mutations may play a relevant oncogenic role. Specifically, diffuse large B-cell malignant lymphoma stands out, with over half of the patients harboring histone driver mutations. Conversely, several other cancer types, including glioblastoma, were depleted for histone mutations. Even though frequent histone driver mutations have been reported in pediatric glioblastoma cases^8,9^, the presence of these mutations is rare among adults^9^.

Previous studies proposed that cancer mutations generally promote oncogenesis by disrupting protein-protein interaction networks^43^. However, the global influence of cancer mutations on epigenetic mechanisms has remained largely unexplored. Nucleosomes function as key hubs in epigenetic signalling pathways, with histone proteins playing central roles in facilitating interactions with DNA, other histones, and NHPs. Based on this, we investigated whether chromatin destabilization at the nucleosomal level and the disruption of epigenetic interactions might be predominant effects of histone mutations. While mutations in specific histone types were statistically associated with PTM sites, histone-histone, and histone-DNA interactions, the strongest enrichment was observed at acidic patch residues in H2A/H2B. The acidic patch is a region that facilitates interactions between nucleosomes^44^ and NHPs, and it was previously suggested that the acidic patch may be a common target for oncogenic histone mutations^4^. We also found that histone mutations (including predicted drivers) were significantly enriched at interfaces with specific NHPs, suggesting that histone mutations may drive cancer development through mechanisms other than direct disruption of nucleosome structures or PTM sites. These mutations likely interfere with the specific functions of histone-binding partners. For example, H4 cancer-specific driver mutations were found to disproportionately accumulate on binding interfaces with 24 different NHPs. This finding was surprising, given H4’s status as a highly conserved histone with no previously documented driver mutations.

Previous work showed that some histone mutations affect DNA-repair pathways and associate with high TMB^25,26^. Therefore, we tried to determine if histone mutations at interaction interfaces with specific NHPs are associated with high TMB. From these analyses, we suggest that disrupting interactions between H4 and TONSL or DAXX may lead to high patient TMB. Indeed, TONSL assists in directing DNA repair upon double-stranded breaks by recognizing H4 proteins and facilitating the deposition of RAD51 on DNA^45,46^. DAXX plays several roles in the cellular DNA damage response, such as promoting p53 activation^47^, regulating RAD51 expression^48^, and depositing H3.3 to newly synthesized DNA after DNA damage^49^. While H3.3 and DAXX perturbations have previously been implicated in cancer^9,50,51^, the effects of H4 cancer mutations on H4-DAXX binding have not been previously reported.

The application of a computational framework integrating genomic and proteomic data allowed us to identify histone mutations and the predominant mechanisms by which they may drive oncogenesis. We propose that perturbing genomic integrity is an emerging mechanism of oncogenesis for histone mutations. We demonstrated that these mutations can disrupt interactions with several histone-binding partners, each having distinct roles in cellular function. Future studies aimed at exploring these mechanisms should control for possible confounding effects of co-occurring mutations and histone gene redundancy. Although most histone mutations are rare, we show that analyzing them based on common structural and functional features is a powerful strategy to illuminate their relationship to cancer.

## Methods

### Identification of Somatic Histone Mutations

Genetic and clinical data were gathered from patients in The Cancer Genome Atlas, Multiple Myeloma Research Foundation CoMMpass, Human Cancer Model Initiative, Count Me In, and Clinical Proteomic Tumor Analysis Consortium programs through the Genomic Data Commons (GDC)^52^. Patients with benign neoplasms were identified based on their primary diagnosis and removed from the dataset. Primary diagnoses follow morphological diagnosis guidelines from the third edition of the International Classification of Diseases for Oncology and can be derived from the clinical data provided by the GDC. WES data were gathered for the neoplasms of cancer patients, in the form of Mutation Annotation Format (MAF) files. All MAF files follow the GDC Ensemble protocol and are readily available for open-access download via the GDC^52^. Patients with multiple MAF files were identified and a union of all variants was used to create a single patient-specific MAF file. For cancer-type specific analyses, patient-level WES data were grouped by patient primary diagnosis. For pan-cancer all patients were grouped together regardless of primary diagnosis.

### Identification of Histone Coding Genes

To identify all human histone somatic mutations in the WES dataset, a comprehensive list of histone genes was compiled by aggregating publicly available data from the Human Genome Organization Gene Nomenclature Committee^17^ and Histone Database 2.0^53^. In total, the list of histone genes includes 96 distinct histone genes, which encode 59 distinct proteins (Supplementary Table S31). Within the list of histone proteins, and throughout the analysis, only full-length proteins that are encoded by the entirety of the protein’s respective coding sequence were considered. Truncated proteins resulting from alternative splicing were not considered. Although some histone genes are tissue-specific, the vast majority of canonical and variant genes are expressed in human normal tissues ^54^. For example, from the Human Protein Atlas, which includes gene expression data from 44 normal human tissue types, 93% of histone genes were expressed in at least one tissue^54^. Based on mass-spectroscopy data, 91% of histone proteins were detected, confirming the existence of their protein form^55^.

### Mutation Driver Status Prediction

To predict the driver status of missense mutations in histone genes, we used the “rank” module of the MutaGene standalone Python package (https://github.com/Panchenko-Lab/mutagene)^28^. MutaGene creates background mutational models (profiles) by aggregating mutation counts for all 96 possible trinucleotide context-dependent point mutations across patients within a given cancer type^27^. To rank mutations by driver status, MutaGene normalizes the observed frequency of the mutation in a given cancer type by an expected frequency derived from the cancer type’s mutational profile. In the case of missense mutations, it considers a penta-nucleotide context around the mutated site. MutaGene reports a B-score, which is the probability of observing a particular mutation in a specific genomic location for a given patient cohort, given its expected frequency. Following mutation ranking, the default MutaGene ranking thresholds were applied to classify mutations as “drivers”, “potential drivers” or “passengers”. As calculations are performed within the scope of a given cancer type, these labels only apply to the individual cancer type and are not transferable across cancer types. Moreover, because mutation rankings are performed at the gene level, driver status labels do not apply to other genes that encode the same protein, as they may vary in their nucleotide sequence. For stringency, only mutations predicted as “drivers” were included in our analyses, whereas “potential drivers” were not.

When performing MutaGene ranking, patients with extremely high TMBs were removed. TMBs were calculated by dividing the total number of mutations in each sample by 30 Mbp, the size of the human exome. Since TMB varies by cancer type, we calculated TMB Z-scores for each patient based on the TMB distribution in the relevant cancer type, and calculated the modified Z-scores for cancer types where TMBs were not normally distributed (Supplementary Note 1). Patients with an absolute Z-score > 3 or an absolute modified Z-score > 3.5 were considered outliers and removed before MutaGene profile creation. Outliers were identified within each cancer type and only cancer types with at least 10 patients were retained for cancer driver mutation status prediction, including the creation of background mutational profiles (Supplementary Table 32).

### Annotation of Histone Interaction Interfaces and Post-Translational Modifications

Histone interactions were extracted from the structural (provided by atomic resolution X-ray crystallography, NMR, and Cryo-EM PDB structures) and chemical cross-linking data of our previously reported human histone interactome (Supplementary Fig. 6A)^33,56^. For PDB structures, an interaction was considered if at least one heavy atom (non-hydrogen) of a histone was within 5 Å of a heavy atom of a binding partner^33^. In the chemical cross-linking data, lysine residues which formed lysine cross-links were considered to form interaction interfaces^33,57^. Interactome data were partitioned into the following types of interaction: histone-histone, histone-DNA, and histone-NHP interactions. Binding partners in the NHP category include proteins that are not histones. A total of 54 histone proteins and 435 binding partners (including DNA, histones, and NHPs) are represented in the interactome. Histone post-translational modifications (PTMs) were gathered from previous data^58^. The list of histone PTMs (Supplementary Table 33) includes PTMs found on most histones within a histone type, as well as PTMs that are unique to individual histone proteins. In total, 818 PTM sites and 20 different chemical modifications were considered across 59 histone proteins. Histone interaction interface, PTMs, and missense mutations were mapped to full-length histone sequences from UniProt^59^.

### Inference of interfaces using multiple sequence alignments

Given that some human histone proteins are not adequately represented in the interactome, empirically derived interactions were supplemented with inferred interactions based on multiple sequence alignments (Supplementary Fig. 6B-E). Multiple sequence alignments were constructed for human histone proteins within the same histone type using Expresso^60^. Alignments included histone tails, H1, and full macroH2A histones. Interaction interfaces observed in individual histone proteins were mapped to aligned positions. These interaction interfaces were then inferred across all histone proteins at the aligned positions that did not report a gap. Positions in the alignment where more than 30% of the aligned proteins showed a gap were not considered eligible for this interface inference process.

### Testing associations of histone mutation occurrence with interfaces and other functionally important sites

Two-tailed Fisher’s exact and binomial tests were performed to test statistical associations between mutation occurrence and functionally important sites (Supplementary Notes 2, 3). To determine if histone mutations were associated with interaction interfaces of a specific type (histone-histone, histone-DNA, or histone-NHP), tests were conducted within the context of an individual histone type (H1, H2A, H2B, H3, H4) and further stratified by one of the three groups: all mutations, mutations observed in more than one patient (recurrent mutations), or mutations restricted to a single patient (non-recurrent mutations). Mutation recurrence was defined at the protein level (see Supplementary Note 4 for detailed information). A Benjamini Hochberg multiple testing correction with a false discovery rate of (FDR) 0.05 was applied to all Fisher’s exact and binomial tests, separately. Adjusted p-values (q-values) less than 0.05 were considered significant. Odds ratios were paired with Fisher’s exact tests to determine effect size, whereas fold-change was calculated for binomial tests (Supplementary Notes 2, 3). All tests were additionally performed using inferred interaction interfaces. Furthermore, additional tests were conducted to determine if histone mutations were associated with inferred H1-DNA interfaces as interaction inference greatly expands the number of H1-proteins considered. Multiple testing correction was repeated as previously described for tests using inferred interaction interfaces.

To determine if histone mutations were associated with PTM sites, tests were conducted within individual histone types. To examine associations between histone mutations and acidic patch residues, tests were conducted considering H2A and H2B proteins collectively, as acidic patch residues occur on both of these histone types. Tests for association at acidic patch residues were conducted by comparing acidic patch residues to all other H2A/H2B residues or to interfacial H2A/H2B residues (not including acidic patch residues). All associations were examined using Fisher’s exact and binomial tests, with multiple testing corrections, significant thresholds, and effect sizes calculated as previously described.

Fisher’s exact and binomial tests were also conducted to determine if histone mutations were associated with interaction interfaces for individual NHPs. Histone proteins were examined collectively rather than being stratified by histone type. Tests were conducted within individual cancer types, in addition to pan-cancer, and were only performed if there were at least ten histone mutations available. Multiple testing corrections, significance thresholds and effect sizes were calculated as previously described. This procedure was repeated when only considering cancer-specific driver mutations.

### Predicting the impact of histone mutations on physical interactions

SAAMBE-3D was used to predict the effects of mutations on histone-protein binding affinity^22^ and SAMPDI-3D on histone-DNA interactions^23^ (ΔΔG, measured in kcal/mol). These machine-learning methods use various features (side chain volume, hydrophobicity, hydrogen bond donor/acceptor, polarity, mutation type, chemical nature of the amino acid residue and others) and are trained on experimental sets of binding affinity changes upon mutations. Both SAAMBE-3D and SAMPDI-3D require experimental structures with atomic resolution to predict ΔΔG. As a result, interactions based exclusively on XLMS data could not be examined. MutaBind2 was used to verify predicted changes in binding affinity since it is more time-consuming and requires structural modelling and minimization (Supplementary Table 30)^24^. SAAMBE-3D and SAMPDI-3D were chosen as the primary methods to predict binding changes given that they are relatively fast in comparison to other methods and perform well.

### Identifying binding partners that have interfaces disproportionately impacted by histone mutations

For each protein binding partner, a Mann-Whitney U test was performed to compare the predicted ΔΔG of histone mutations at interfaces for that partner to the predicted ΔΔG of mutations at interfaces for all other partners. Rank-biserial correlation coefficients were calculated alongside each Mann-Whitney U test. As several structures may be used to obtain ΔΔG estimates for a given mutation-partner pair, the mean ΔΔG estimate across all structures was used for this analysis. Tests were only conducted if there were at least 10 unique mutations with a ΔΔG prediction available for interactions with a given protein binding partner and at least 10 mutations with a ΔΔG prediction available for interactions with other protein binding partners. Tests were conducted for pan-cancer and each cancer type, only considering mutations that were observed in the given cancer type. A Benjamini-Hochberg correction was applied throughout all tests, at an FDR of 0.05. Tests reporting a q-value less than 0.05 were considered significant.

### Comparisons of Tumor Mutational Burden

Kruskal Wallis tests were conducted to determine if there was a significant difference in TMB between patients without histone mutations, patients with histone mutations not at NHP interfaces, and patients with histone mutations at NHP interfaces. This was done for pan-cancer and individual cancer types. Only cancer types with at least ten patients in each of the groups above were examined. Two kinds of Kruskal Wallis tests were used (see Supplementary Note 5). Post-hoc Dunn tests were conducted to determine pairwise differences between groups in instances where both Kruskal-Wallis tests reported a significant difference (q < 0.05, Benjamini-Hochberg, FDR = 0.05). A rank-biserial correlation coefficient was calculated to accompany each pairwise comparison. This analysis was repeated after removing patients with *POLE* mutations.

For each NHP type, patients with a histone mutation at an interface for a specific NHP were identified and placed in an “NHP group”. For each NHP group, Mann-Whitney U tests were conducted to determine if the TMB of patients in the NHP group differed significantly from patients without histone mutations, patients with histone mutations not on NHP interfaces, and patients with histone mutations at interfaces for other NHPs. For each pairwise comparison, a rank-biserial correlation coefficient was calculated. Two types of Mann-Whitney U tests were conducted (see Supplementary Note 5). For either test type, a significant difference was defined as a q-value less than 0.05 after multiple testing corrections with the Benjamini-Hochberg method (FDR = 0.05). Tests were only conducted if both groups had at least ten patients. Tests were performed for pan-cancer and individual cancer types. This analysis was also performed after removing patients with *POLE* mutations.

## Acknowledgements

The authors would like to thank Dr. Alexey Shaytan of the Lomonosov Moscow State University for generously sharing data related to histone gene nomenclature. D.E., D.O., S.L. and A.R.P. were supported by the Department of Pathology and Molecular Medicine, Queen’s University, Canada. Y. S. was supported by the Department of Biology and Molecular Sciences, Queen’s University, Canada. A.R.P. is the recipient of a Senior Canada Research Chair in Computational Biology and Biophysics and a Senior Investigator Award from the Ontario Institute of Cancer Research, Canada. A.R.P. also acknowledges the support of the Natural Sciences and Engineering Research Council of Canada (NSERC) (No. RGPIN/02972-2021). M.J.A. is supported by a Faculty of Arts and Science infrastructure and a Research Initiation grant. This research is supported by New Frontier in Research Fund Exploration (NFRFE-2021-00880) and Cancer Research Society Operation grants (1056783) to A.R.P and M.J.A. D.L. was supported by the Intramural Research Program of the National Library of Medicine, National Institutes of Health. The views expressed in the publication are the views of the authors and do not necessarily reflect those of the Government of Ontario. Y.P. is supported by the National Natural Science Foundation of China (No. 12205112), Natural Science Foundation of Wuhan (No.2024040801020302) and self-determined research funds of CCNU from the colleges’ basic research and operation of MOE.

